# PepMCP: A Graph-Based Membrane Contact Probability Predictor for Membrane-Lytic Antimicrobial Peptides

**DOI:** 10.64898/2026.02.01.703163

**Authors:** Ruihan Dong, Tadsanee Awang, Qiushi Cao, Kai Kang, Lei Wang, Zefeng Zhu, Chen Song

**Affiliations:** Center for Quantitative Biology, Peking-Tsinghua Center for Life Sciences, Academy for Advanced Interdisciplinary Studies, Peking University, Beijing 100871, China; Peking University–Tsinghua University–National Institute of Biological Sciences Joint Graduate Program, Academy for Advanced Interdisciplinary Studies, Peking University, Beijing 100871, China

**Keywords:** Antimicrobial peptide, Membrane contact probability, Graph neural network, Membrane-lytic mechanism

## Abstract

**Motivation:** The membrane-lytic mechanism of antimicrobial peptides (AMPs) is often overlooked during their in silico discovery process, largely due to the lack of a suitable metric for the membrane-binding propensity of peptides. Previously, we proposed a characteristic called membrane contact probability (MCP) and applied it to the identification of membrane proteins and membrane-lytic AMPs. However, previous MCP predictors were not trained on short peptides targeting bacterial membranes, which may result in unsatisfactory performance for peptide studies.

**Results:** In this study, we present PepMCP, a peptide-tailored model for predicting MCP values of short peptides. We collected more than 500 membrane-lytic AMPs from the literature, conducted coarse-grained molecular dynamics (MD) simulations for these AMPs, and extracted their residue MCP labels from MD trajectories to train PepMCP. PepMCP employs the GraphSAGE framework to address this node regression task, encoding each peptide sequence as a graph with 4-hop edges. PepMCP achieved a Pearson correlation coefficient of 0. 883 and an RMSE of 0. 123 on th e node-level test set. It can recognize membrane-lytic AMPs with the predicted MCP values for each sequence, thereby facilitating mechanism-driven AMP discovery. Additionally, we provide a database, MemAMPdb, which includes the membrane-lytic AMPs, as well as the PepMCP web server for easy access.

**Availability and Implementation:** The code and data are available at https://github.com/ComputBiophys/PepMCP.

**Supplementary Information:** Supplementary data are available online.

## Introduction

Antimicrobial peptides (AMPs) are short peptides widely present in the innate immune system of various organisms. AMPs are known for their primary mechanisms of disrupting membranes, which provide them with broad-spectrum antimicrobial abilities and reduce the likelihood of causing drug-resistance (Júnior et al., 2025). Recently, machine learning techniques have facilitated the discovery of new AMPs by screening hundreds of metagenomes (Ma et al., 2022; Santos- Júnior et al., 2024), proteomes (Li et al., 2025c), and theoretical sequence spaces (Huang et al., 2023; Szymczak et al., 2025). However, the membrane-lytic mechanism of AMPs is usually not considered in their computational procedures due to the lack of a metric to characterize the membrane-lytic propensity of peptides.

Conventional physiochemical properties, such as hydrophobicity, hydrophobic moment, and the Boman index, provide a general description of sequences; however, they are coarse metrics and cannot serve as indicators for membrane-lytic AMPs (Santos-Júnior et al., 2024). There are some structure-dependent tools, such as DREAMM for predicting protein-membrane interfaces (Chatzigoulas and Cournia, 2022; Paranou et al., 2024) and PPM for predicting the spatial orientation of proteins in membranes (Lomize et al., 2022). However, these tools were not primarily developed for short peptides. PMIpred (van Hilten et al., 2024) trained a neural network model on binding free energies calculated from molecular dynamics (MD) simulations, but it only used fixed-length peptides (24 residues) and focused on recognizing curvature-sensing peptides, which might not be applicable to membrane-lytic AMPs of different lengths.

Previously, we proposed a characteristic called membrane contact probability (MCP) for studying the structure and function of membrane proteins. MCP is defined as the likelihood of each residue in a protein sequence to be in direct contact with the hydrophobic cores of membranes (Wang et al., 2022). MCP can be extracted from MD simulations by calculating the fraction-of-time probability of the residue *α*-carbon being within 6 Åof the lipid acyl chain carbon atoms. Given the time-consuming nature of running MD simulations, we developed deep learning-based MCP predictors and applied them in contact map prediction (Wang et al., 2022), membrane protein screening and design (Wang et al., 2025a; Li et al., 2025a), and studying mechanosensitive protein dynamics (Han et al., 2025). In particular, we utilized MCP to discover novel membrane-lytic AMPs from human and frog metaproteomes. We constructed a pipeline incorporating the prediction of MCP, helical propensity, and anti-parallel dimerization, and we successfully discovered seven membrane-lytic AMPs (Li et al., 2025b).

Even though MCP has shown potential in studying membrane-lytic AMPs, some limitations exist with previous MCP predictors. First, they were primarily developed for membrane proteins, and their training data lacked peptides. The minimal sequence length in their training set was restricted to 26 amino acids, which affected the prediction accuracy for short peptides. Second, the MCP labels were derived from the MemProtMD database (Newport et al., 2018), where the membranes in MD simulations were composed of pure 1,2-dihexadecanoyl-rac-glycero-3-phosphocholine (DPPC) lipids, mimicking mammalian membranes rather than bacterial membranes. Although some membrane proteins exhibited similar MCP distributions with phosphatidylcholine (PC) or phosphatidylethanolamine (PE)/phosphatidylglycerol (PG) lipids (Wang et al., 2022), many AMPs demonstrated different behaviors and exhibited membrane selectivity (Suarez-Leston et al., 2022; Dong et al., 2025). Therefore, it is crucial to develop a peptide-specific MCP predictor for bacterial membranes, contributing to the discovery of membrane-lytic AMPs.

In this study, we built a peptide-tailored MCP predictor, PepMCP, based on the graph sample and aggregate (GraphSAGE) model. To train the PepMCP model, we collected a high-quality dataset that contained 516 membrane-lytic AMPs and conducted coarse-grained (CG) MD simulations to calculate their MCP values while interacting with bacterial membranes (Fig. 1a). PepMCP encoded a peptide as a graph with 4-hop edges and ESM C node embeddings to capture spatial information without requiring peptide structures (framework in Fig. 1b). PepMCP predicted the residue MCP values of each node and achieved a Pearson correlation coefficient of 0.883 and a root mean square error (RMSE) of 0.123 on the node-level split test set. We demonstrate that PepMCP can not only predict the MCP patterns in peptides, but also be utilized to recognize membrane-lytic AMPs from soluble peptides using sequence average MCP values.

**Fig. 1.**
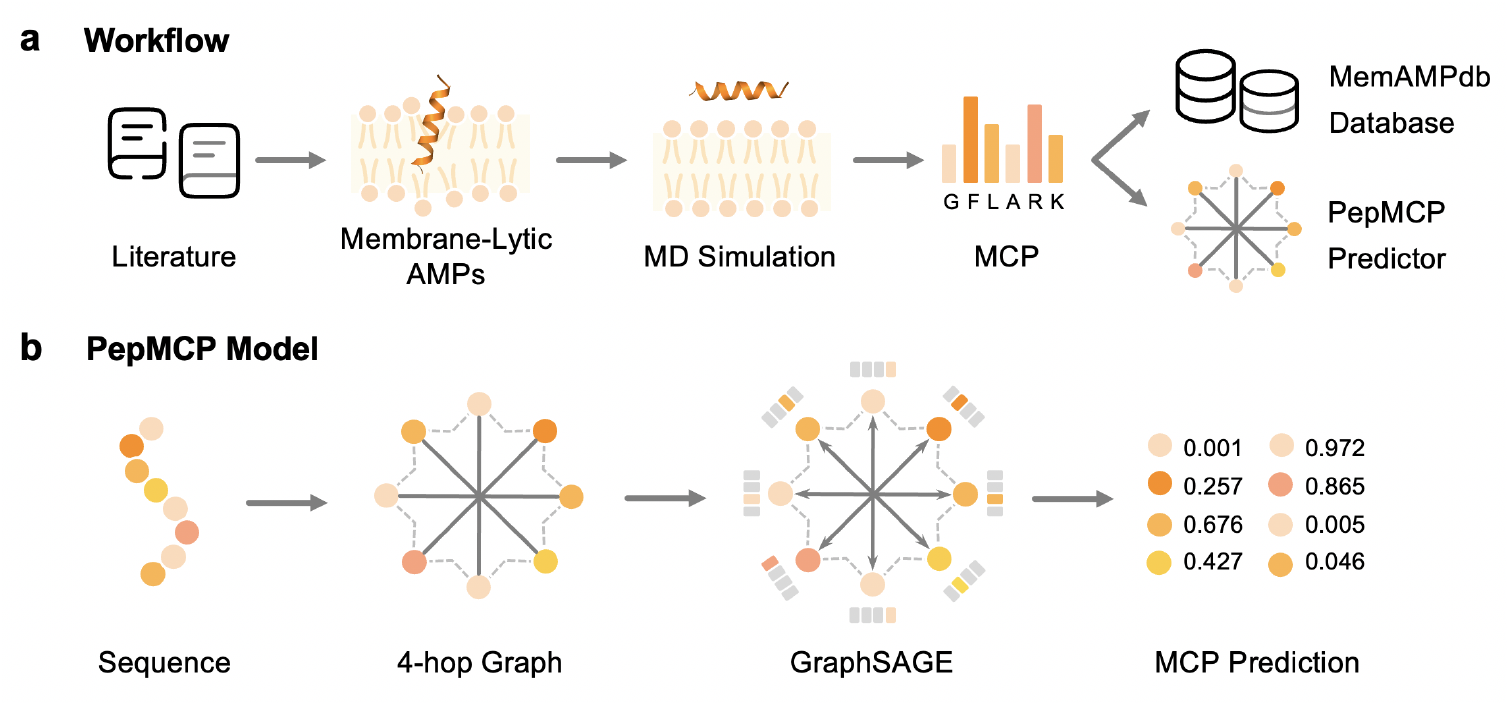
Overview of the study. **a**. Workflow of PepMCP, including collecting the membrane-lytic AMPs from literature, running coarse-grained MD simulations, calculating the membrane contact probability (MCP) values of each residue, training the PepMCP predictor, and building the MemAMPdb database. **b**. Framework of the PepMCP model. PepMCP receives peptide sequences as input, encodes them as 4-hop graphs, and utilizes the GraphSAGE model to accomplish the node-level regression task of predicting MCP values. The dashed lines in the octagram represent the sequential order of residues in a peptide, while the solid lines represent 4-hop connected edges.

## Materials and Methods

### Data Collection

A membrane-lytic AMP dataset was manually curated from literature. Using the keyword ‘antimicrobial peptide’, the PubMed database was searched for publications reporting experimentally validated membrane-lytic AMPs (as of November 2024). The membrane-lytic mechanisms of AMPs were confirmed with the following approaches: (1) membrane permeability assays, in which membrane integrity–sensitive fluorescent dyes were incubated with bacteria and peptides. Propidium iodide (PI) was used to indicate inner membrane integrity. N-phenyl-1-naphthylamine (NPN) was used to indicate outer membrane permeabilization. 3,3’-dipropylthiadicarbocyanine iodide (DiSC_3_(5)) was used to assess cytoplasmic membrane depolarization (Ma et al., 2022); (2) liposome leakage assays, in which bacterial membrane–mimicking liposomes encapsulating fluorescent dyes were used to evaluate AMP-induced membrane disruption (Ambroggio et al., 2005); (3) scanning electron microscopy (SEM) and transmission electron microscopy (TEM), to examine morphological alterations of bacterial membranes; and (4) fluorescence microscopy and flow cytometry, combined with fluorescent dyes such as PI or SYTOX green, to detect changes in membrane integrity, where increased fluorescence generally indicates membrane disruption (Buck et al., 2019).

During this collection process, AMP sequences with complex chemical modifications, such as cyclization, fatty acid conjugation, or non-canonical amino acids, were excluded. Sequences with lengths ranging from 10 to 51 amino acids were retained. Redundant sequences were removed using CD-HIT (Li et al., 2001) at a threshold of 90%. Ultimately, the membrane-lytic AMP dataset comprised 516 sequences, which were used as the positive set with MCP labels obtained from our MD simulations.

For the negative set, soluble peptides were collected from the Protein Data Bank (as of May 2020) following a similar procedure to that of the MCP predictor (Wang et al., 2022). The sequence lengths ranged from 10 to 51 residues. Peptides with ‘antimicrobial’, ‘antibiotic’, or related annotations were discarded. A total of 1307 sequences remained after redundancy removal using CD-HIT with a threshold of 70%. Then, 516 peptides were randomly selected as the negative set, and their residues were labeled with zero.

The training, validation, and testing sets were partitioned using both the residue-level and sequence-level split approaches. On the residue level, 20% of nodes were retained for testing, while the remaining 80% of nodes were divided into training and validation sets through a 5-fold cross-validation approach for each sequence. Finally, 24,636 nodes were included in the training and validation sets, while 5,663 nodes were included in the test set. On the sequence level, 206 sequences (20% of the total) and all their constituent nodes were designated for testing. The remaining 826 sequences were used for 5-fold cross-validation.

### MD Simulation

The structures of 516 membrane-lytic AMPs were predicted using ColabFold 1.5.5 (Jumper et al., 2021; Mirdita et al., 2022) and mapped to a CG representation via the *martinize*.*py* script (de Jong et al., 2013). An elastic network was adopted to maintain the secondary structures with a force constant of 500 *kJ mol*^*−*1^ *nm*^*−*2^ and an interaction cut-off range between 0.5 nm and 0.9 nm (Monticelli et al., 2008; Periole et al., 2009;Poma et al., 2017). The membrane bilayers were constructed using the CHARMM-GUI Martini Maker (Wu et al., 2014; Qi et al., 2015), consisting of 1-palmitoyl-2-oleoyl-sn-glycero-3-phosphoethanolamine (POPE) and 1-palmitoyl-2-oleoyl-sn-glycero-3-phosphoglycerol (POPG) in a 3:1 molar ratio. To optimize computational efficiency, systems were scaled into three sizes based on the longitudinal dimension of the peptide (Fig. S1a): Small (*≤* 3 nm, 40 lipids), Medium (3–5 nm, 69 lipids), and Large (*>* 5 nm, 168 lipids). Each lipid bilayer underwent a 100-ns equilibration to achieve phase stability. Peptides were then positioned in the aqueous phase, parallel to the membrane surface at an initial distance of 15–20 Å. All systems were solvated with standard Martini water beads and ionized with 150 mM NaCl for neutralization.

All MD simulations were performed using GROMACS 2022.5 (Abraham et al., 2015) with the Martini 2.2 force field (de Jong et al., 2013). A 5000-step energy minimization was conducted using the steepest descent algorithm. Systems were equilibrated for 1 ns in the NVT ensemble, followed by a 50 ns NPT equilibration, during which harmonic positional restraints were applied to the peptide backbone beads. For production simulations, unrestrained MD trajectories were produced for 2–5 *µs* per system, with a time step of 20 fs. The temperature was kept at 310 K using the v-rescale thermostat (Bussi et al., 2007) and the pressure at 1 bar using the Parrinello-Rahman barostat with semi-isotropic coupling (Parrinello and Rahman, 1981). Non-bonded interactions were calculated using a cut-off of 1.2 nm for both van der Waals and electrostatic forces. The latter was treated using the reaction-field method with a dielectric constant (*ϵ*_*rf*_) of 15.

### MCP Calculation

The membrane-binding property of the 516 independent trajectories was evaluated using the minimum distance between the protein backbone and the lipid phosphate groups (BB–PO_4_). Of the 516 membrane-lytic AMPs, 514 showed stable membrane binding over the final 1 *µs* of MD trajectories (Fig. S1b, S2a and b), while two outliers did not exhibit stable interactions with membranes over the 5 *µs* trajectories in our simulations (Fig. S2c and d). This suggests that the majority of the AMPs in the positive dataset are membrane-interacting peptides, and the MCP values obtained from MD simulations can be used to characterize the membrane-binding features of these AMPs.

The MCP between individual peptide residues and the lipid bilayer was calculated using the final 1 *µs* of MD trajectories. A contact was defined based on a spatial proximity threshold of 6.0 Å between a residue’s backbone (BB) bead and any bead within the hydrophobic lipid tail (C1A, C1B, D2A, C2B, C3A, C3B, C4A, and C4B) of both POPE and POPG (Wang et al., 2022). MCP values were calculated using the MDAnalysis library (Michaud-Agrawal et al., 2011).

### PepMCP Model

#### Model Framework

Each peptide sequence was encoded as a graph *G* =*< V, E >*. The nodes (*V* = *{v*_0_, *v*_1_, …, *v*_*n*_*}*) represented *n* residues in the sequence, and the node features (*H* ∈ ℝ^*m×n*^) were derived from a protein language model (ESM C 300M (Hayes et al., 2025), where the feature dimension *m* = 960). The edges were encoded using the 4-hop approach, which meant that an edge *e* connects *v*_*i*_ and *v*_*i*+4_. This edge encoding effectively extracted the peptide structural information, as most membrane-lytic AMPs are *α*-helix.

The GraphSAGE (Hamilton et al., 2017) method was adopted as our model to process peptide graphs. GraphSAGE is an inductive approach that includes three steps: neighborhood sampling, feature aggregation, and label prediction. Using an aggregate function *aggregate*(*·*), the feature of node *v* ∈ *V* at layer (*k* + 1) would be:

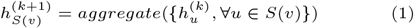

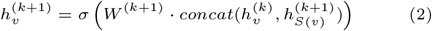

where *S*(*v*) is the neighborhood node set of *v, σ*(*·*) is the activation function ReLU, *W* is a trainable weight matrix, and *concat*(*·*) is the concatenation operation.

The pooling aggregator was used in PepMCP. It conducted an element-wise max pooling after a linear transformation of the neighborhood node features:

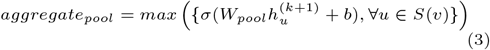

Three layers were included here to compress the 960-dimension feature to 512-dimension, and predicted the 1-dimension label. The outputs were transformed into probability values using the sigmoid function.

#### Model Implementation

PepMCP was developed using the PyTorch and DGL libraries (Wang et al., 2019). The loss function of PepMCP was the mean squared error. The optimizer used was Adam, with a learning rate of 0.0001. The batch size for the graph was 4. The training epoch was set to 20, and early stopping was adopted when the model showed no improvement in the validation loss for 10 epochs.

Four regression metrics were used to evaluate the model performances, including the Spearman’s rank correlation coefficient (Spearman), the Pearson correlation coefficient (Pearson), the coefficient of determination (*R*^2^), and the root mean square error (RMSE).

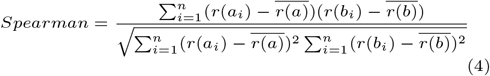

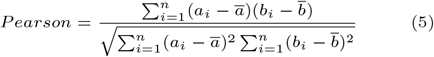

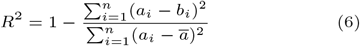

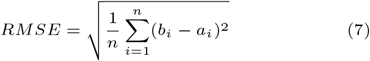

where *a*_*i*_ and *b*_*i*_ denote the actual and predicted values of the sequence *i*, respectively. *r*(*·*) calculates the ranking number.

### Application on Independent Membrane-Lytic AMPs

An external test set was curated to demonstrate the application of PepMCP on membrane-lytic AMPs. A total of 34 novel membrane-lytic AMPs were collected from the literature published in 2024-2025, excluding those that overlapped with the sequences in the training set. These membrane-lytic AMPs were also confirmed to have at least one of the aforementioned experimental proofs. Similarly, coarse-grained MD simulations were conducted using AlphaFold structures of these 34 peptides. The residue-level MCP values were then calculated as the ground truth labels. A collection of soluble peptides was obtained from the PDB (after May 2020) with lengths ranging from 10 to 51 residues, excluding those associated with membrane-related or antibiotic-related annotations. Consequently, 46 soluble peptides remained and were assigned zero labels. This external test set includes 80 peptides and 2133 residue nodes.

## Results

### Membrane-Lytic AMPs and their MCP Values

Since current AMP databases often lack clear and verifiable annotations of mechanisms, we curated a high-quality membrane-lytic AMP dataset containing 516 AMPs with experimentally validated mechanisms. Most of the AMPs originated from natural sources, while some were generated by deep learning models. Then we utilized MD simulations to investigate the peptide-membrane interactions and calculate the MCP values of each residue. Considering that not all AMPs had solved 3D structures, we used their AlphaFold-predicted structures in MD simulations. 88% of the structures had an average pLDDT *>* 70 (Fig. S3a). Most membrane-lytic AMPs were *α*-helix, with a few *β*-sheet, random coil, or *αβ* secondary structures (Fig. 2a). At the final frame of simulations, these peptides could stably attach to the surface of bacterial membranes (Fig. S1b), often with one side contacting the lipid tails and exhibiting larger MCP values. Fig. 2b shows the MCP frequency distributions of 20 canonical amino acids, where the amino acid orders follow the Eisenberg hydrophobicity scale. Hydrophobic residues such as isoleucine (I), phenylalanine (F), valine (V), and leucine (L) had a higher proportion of MCP values greater than 0.5 and tended to be in contact with the lipid tail via hydrophobic interactions. Alanine (A) and glycine (G) also showed a certain proportion of MCP values close to one, due to their simple side chains. Most MCPs of polar amino acids are close to zero.

**Fig. 2.**
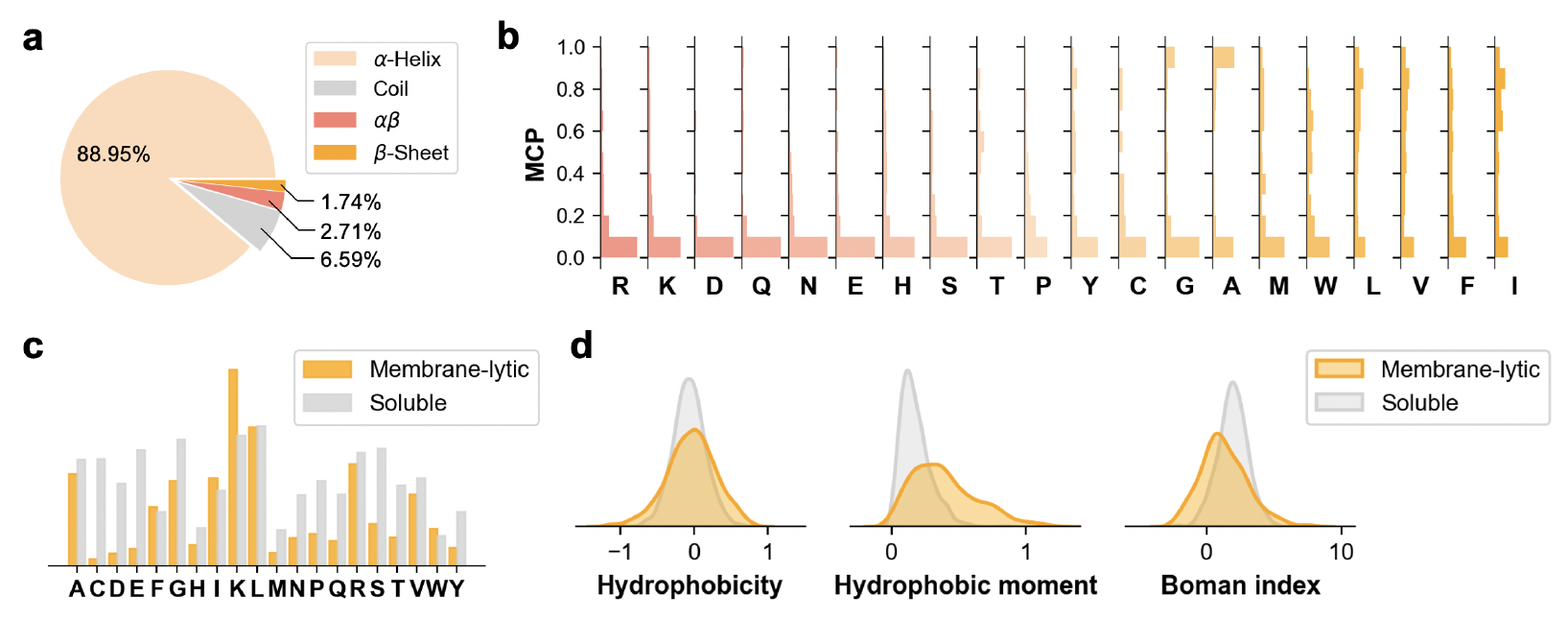
Dataset characterization. **a**. Secondary structures of membrane-lytic AMPs as predicted by AlphaFold2. **b**. Frequency distribution of MCP values across 20 amino acids. The maximum x-axis value (frequency) of each subplot is 0.6. The order of amino acids follows the hydrophobicity. **c**. Amino acid frequency of membrane-lytic AMPs and soluble peptides. **d**. Physicochemical properties (hydrophobicity, hydrophobic moment, and Boman index) of membrane-lytic AMPs and soluble peptides. Eisenberg’s scale was used for all hydrophobicity measurements.

We also included soluble peptides as negative samples to improve the generalization capability of our proposed model. The amino acid frequency of membrane-lytic AMPs and soluble peptides differed, as the former contained a higher frequency of positively-charged lysine (K) and a lower frequency of negatively-charged aspartic acid (D) and glutamic acid (E), which is a characteristic feature of AMPs (Fig. 2c). We compared some physicochemical properties of membrane-lytic AMPs and soluble peptides in Fig. 2d. Membrane-lytic AMPs exhibited a wider hydrophobicity distribution and larger hydrophobic moments. They also showed a relatively smaller Boman index, indicating a propensity for membrane interaction (Boman, 2003). However, there were still overlapping areas between the two peptide sets, so these properties could not be used to distinguish all the membrane-lytic AMPs from soluble peptides, making it essential to develop other indicators.

### PepMCP is a Peptide-Tailored MCP Predictor

Based on the membrane-lytic AMPs and their MCP values, we developed the PepMCP model, focusing on these short peptides. The lengths of all the sequences in the training set ranged from 10 to 51 amino acids. We encoded the peptide sequences using graphs, with each residue representing a graph node. We designed 4-hop edges to capture the spatial interactions of peptides without introducing their 3D structures, by connecting the *i*^*th*^ node with the (*i* + 4)^*th*^ node (Fig. 1b). We implicitly encoded the sequential information of peptide sequences in their node embedding from the ESM C language model. We utilized the inductive GraphSAGE model to process the features, which aggregated the features on the local subgraph and produced new features for the unseen nodes.

We compared the performance of the PepMCP model with that of two previous MCP predictors. One was a special version of the MCP predictor, which used the SPIDER3-Single (Heffernan et al., 2018) features to replace multiple sequence alignments (MSAs), as short peptides often lack abundant MSAs. We utilized this predictor to screen novel membrane-lytic AMPs with dimerization metrics and reached a moderate success rate of 39% (Li et al., 2025b). We referred to this model as the ‘base’ model in the following comparisons. The other model was ProtRAP-LM, which used ESM-2 to encode protein inputs and could predict their relative accessibility simultaneously (Wang et al., 2025a). The minimum sequence length in their training set was 26, as their predictive goal was not primarily short peptides, which resulted in limited predictive ability on our test set (Table. 1). On the residue-level split test set, the base model and ProtRAP-LM could only obtain Pearson correlation coefficients of 0.2186 and 0.1133, respectively. The base model performed slightly better when using the sequence-level split, achieving a Spearman of 0.2069 and a Pearson of 0.2592, while ProtRAP-LM performed worse. In contrast, PepMCP model significantly outperformed the two models, achieving a Spearman correlation of 0.7963 and a Pearson correlation of 0.8825 for the residue level split set (Table. 1). For the sequence-level split, the performance is slightly lower but still achieves correlations above 0.70. Therefore, PepMCP is a customized and capable model for predicting peptide-specific MCP values.

**Table 1.**
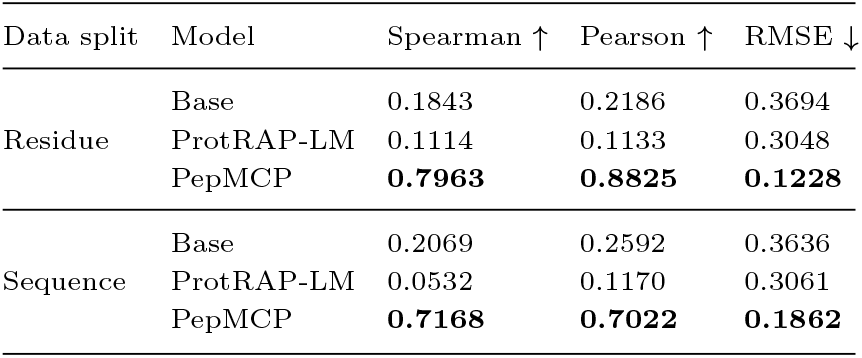
Performance of PepMCP, ProtRAP-LM, and the base model on the test set. Bold values indicate the best performance for the corresponding metric.

| Data split | Model | Spearman ↑ | Pearson ↑ | RMSE ↓ |
| --- | --- | --- | --- | --- |
| Residue | Base | 0.1843 | 0.2186 | 0.3694 |
|  | ProtRAP-LM | 0.1114 | 0.1133 | 0.3048 |
|  | PepMCP | **0.7963** | **0.8825** | **0.1228** |
| Sequence | Base | 0.2069 | 0.2592 | 0.3636 |
|  | ProtRAP-LM | 0.0532 | 0.1170 | 0.3061 |
|  | PepMCP | **0.7168** | **0.7022** | **0.1862** |

### Comparison of Different Architectures

We compared the performances of four graph neural network variants, including the graph convolutional network (GCN) (Kipf and Welling, 2016), GraphSAGE (Hamilton et al., 2017), graph attention network (GAT) (Veličković et al., 2018), and topology adaptive graph convolutional network (TAGCN) (Du et al., 2018). Fig. 3a showed the results of 5-fold cross-validation. The GraphSAGE model achieved the best performance among the four types of GNNs, with a Spearman coefficient of 0.8016 *±* 0.0029, a Pearson correlation coefficient of 0.8762 *±* 0.0060, an *R*^2^ of 0.7664 *±* 0.0107, and an RMSE of 0.1258 *±* 0.0029. The inductive aggregation of GraphSAGE model effectively captured the information of peptide sequences. Regarding the other models, TAGCN also had a comparable Spearman of 0.8016 *±* 0.0039; however, it could not be compared to GraphSAGE on the other three metrics. The aggregator was a crucial component of the GraphSAGE model, so we tested four types of aggregators (Fig. 3b). The max pooling aggregator (‘pool’) demonstrated the best performance across the four regression metrics.

**Fig. 3.**
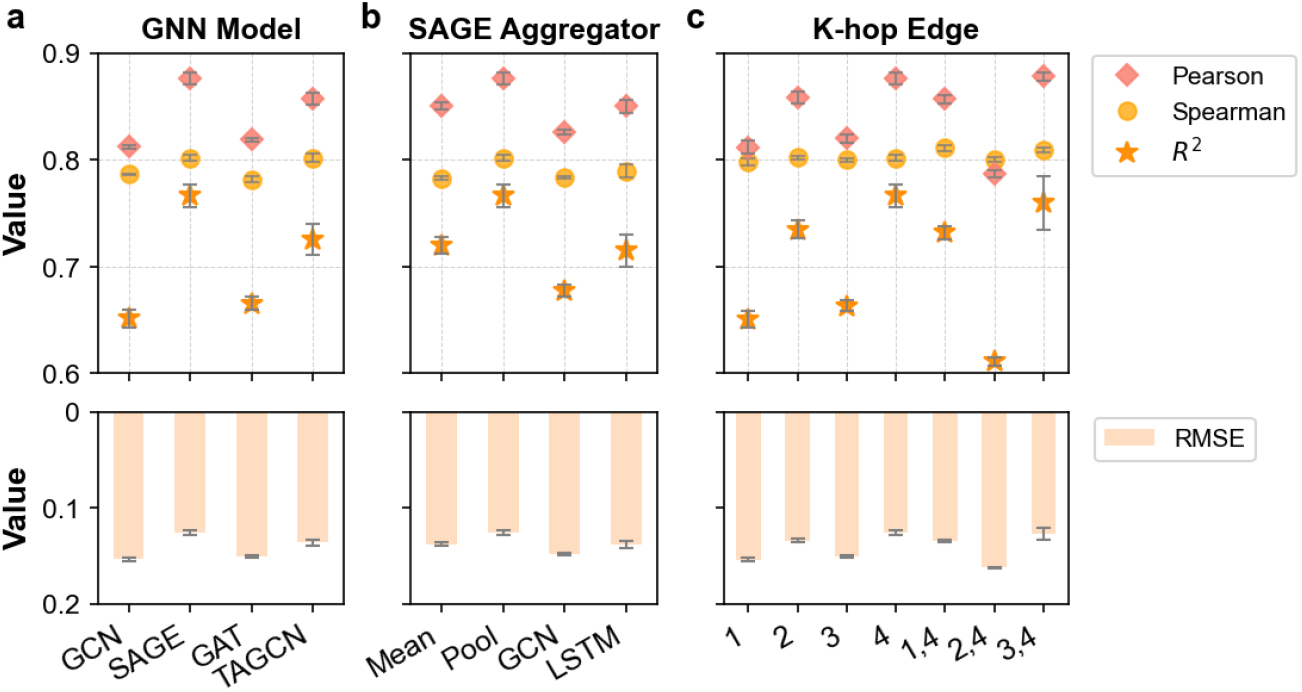
Comparison of different model architectures. **a**. Graph neural network (GNN) variants. **b**. Aggregator of GraphSAGE (abbreviated as SAGE in the figure). **c**. K-hop edge for encoding sequence graphs. Scatter plots show Spearman, Pearson, and *R*^2^ (higher is better), while bar plots show RMSE values (lower is better). All the results are average values on the test set using 5-fold cross validation models, and the error bars represent the standard errors.

Meanwhile, we compared different edge encoding approaches in PepMCP. We tested k-hop edges (connecting edges between nodes *v*_*i*_ and *v*_*i*+*k*_) for k = 1, 2, 3, or 4 (Fig. 3c). Overall, the best performance was observed when k = 4. We suggest that this is related to the secondary structure of peptides, as almost 90% of them were *α*-helix. There are approximately 3.6 residues per turn in the helix, so the *i*^*th*^ and (*i* + 4)^*th*^ residues are spatially adjacent. Therefore, encoding peptide graphs with 4-hop edges is a simple yet effective method to process residue-level predictions. The sequential graph (k = 1) contributed to the second-to-last worst performance, indicating that feature aggregation of the residues along the sequence provided limited useful connections. We also tested double combinations of k-hop edges, encoding k = 4 edges along with another k-hop edge where k = 1, 2, or 3. Even though some of them exhibited slightly higher metrics compared to k = 4, for instance, k = 1, 4 had a Spearman of 0.8110 *±* 0.0026, and k = 3, 4 had a Pearson of 0.8779 *±* 0.0044, they did not demonstrate superior performance compared to k = 4 in terms of *R*^2^ and RMSE values.

For the node features, we compared three lightweight protein language models, Ankh (Elnaggar et al., 2023), ESM C (300M and 600M) (Hayes et al., 2025), and Profluent-E1 (150M, 300M, and 600M) (Jain et al., 2025). Table 2 presents the results of 5-fold cross validation, where ESM C 300M achieved the best performance. Interestingly, the 150M and 300M versions of Profluent-E1 displayed large standard deviations because they were unable to handle one or two folds out of the 5-fold cross validation sets. In brief, the final PepMCP model utilized the GraphSAGE architecture with the pool aggregator, encoded edges in a 4-hop approach, and extracted node features from ESM C 300M.

**Table 2.** Performance using node features from different language models on the test set. Average values *±* standard deviations from 5-fold cross validation were reported. Bold values represent the best performance for that metric.

| Node Feature | Param. | Spearman ↑ | Pearson ↑ | $R^2$ ↑ | RMSE ↓ |
| --- | --- | --- | --- | --- | --- |
| ESM C | 300M | **$0.8016 \pm 0.0029$** | $0.8762 \pm 0.0060$ | **$0.7664 \pm 0.0107$** | **$0.1258 \pm 0.0029$** |
| ESM C | 600M | $0.8000 \pm 0.0039$ | $0.8754 \pm 0.0020$ | $0.7603 \pm 0.0081$ | $0.1274 \pm 0.0021$ |
| Ankh-base | 450M | $0.7956 \pm 0.0038$ | **$0.8775 \pm 0.0037$** | $0.7590 \pm 0.0211$ | $0.1277 \pm 0.0054$ |
| Profluent-E1 | 150M | $0.6812 \pm 0.1191$ | $0.5475 \pm 0.3837$ | $0.3455 \pm 0.4736$ | $0.1964 \pm 0.0760$ |
| Profluent-E1 | 300M | $0.7569 \pm 0.0730$ | $0.7169 \pm 0.3126$ | $0.5615 \pm 0.3976$ | $0.1597 \pm 0.0647$ |
| Profluent-E1 | 600M | $0.7825 \pm 0.0079$ | $0.8753 \pm 0.0028$ | $0.7586 \pm 0.0084$ | $0.1279 \pm 0.0022$ |

### Recognizing Membrane-Lytic AMPs with PepMCP

We prepared an external test set that included 34 new membrane-lytic AMPs and 46 soluble non-AMPs. The average similarity between these sequences and all the 1032 peptides in the training set was 39.7%, as calculated by the local Smith-Waterman alignment. Although PepMCP was trained on residue-level split data, it performed well in predicting the MCP values for all the residues of unseen sequences, and achieved a Pearson of 0.8226 on this external test set (Fig. 3). Similarly, PepMCP significantly outperformed the previous ProtRAP-LM and base models. In addition, we used the average MCP prediction for all residues in a peptide as a sequence-level prediction. In this way, we evaluated PepMCP’s ability to distinguish membrane-lytic AMPs from other peptides using classification metrics. Following the previous threshold (Li et al., 2025b), we regarded a peptide as positive if its average MCP was greater than 0.2. Otherwise, the peptide was predicted to be negative. Table 3 also shows the classification results on this external test set, considering membrane-lytic AMPs as positive samples and soluble non-AMPs as negative samples. PepMCP achieved AUC, accuracy, and precision scores of over 0.9. Although ProtRAP-LM and the base model also obtained AUC values of over 0.7, they were not comparable to PepMCP on other metrics, especially precision. This result demonstrates the effectiveness of PepMCP in distinguishing membrane-lytic AMPs from soluble peptides.

**Table 3.** Results of both classification and regression tasks on the external test set. Bold values indicate the best performance for that metric.

| Task | Metric | PepMCP | ProtRAP-LM | Base |
| --- | --- | --- | --- | --- |
| Regression | Spearman | **0.7182** | 0.3723 | 0.2587 |
|  | Pearson | **0.8226** | 0.0706 | 0.2868 |
| | $R^2$ | **0.6750** | -0.3416 | -0.8361 |
|  | RMSE | **0.1531** | 0.3112 | 0.3640 |
| Classification | AUC | **0.9277** | 0.7168 | 0.7008 |
|  | AUPR | **0.8606** | 0.5502 | 0.5991 |
|  | Accuracy | **0.9000** | 0.5875 | 0.4875 |
|  | Precision | **0.9333** | 0.5294 | 0.4493 |
|  | Recall | 0.8235 | 0.2647 | **0.9118** |
|  | F1 | **0.8750** | 0.3529 | 0.6019 |

We presented four cases from the external test set in Fig. 4, two of which were membrane-lytic AMPs (Fig. 4a) and two were soluble non-AMPs (Fig. 4b). PepMCP effectively captured the zigzag MCP patterns of this amphipathic *α*-helix peptide p11, which was mined from human gut microbe metagenomes (Li et al., 2025c). MCP coloring in its AlphaFold structure showed that its hydrophobic side had a high tendency to contact the lipid bilayer. A short AMP m AMP76 (9 residues) was screened from UniProtKB by AMPSorter (Wang et al., 2025b). PepMCP also predicted well on this random coil, with only slight errors for two tryptophan (W) residues. Regarding the negative samples, PepMCP predicted MCP values that were very close to zero for a glucagon analog peptide (PDB ID: 6PHO) and a Z0 domain from the transcription repressor BCL11A (PDB ID: 9BV0) (Harris et al., 2025). In Fig. 4b, we zoomed in on the view of MCP values and found that there were still periodic patterns every four residues in the predictions of *α*-helix 6PHO. This phenomenon also existed in the C-terminal *α*-helix part of 9BV0, but did not appear in the *β*-sheet part.

**Fig. 4.**
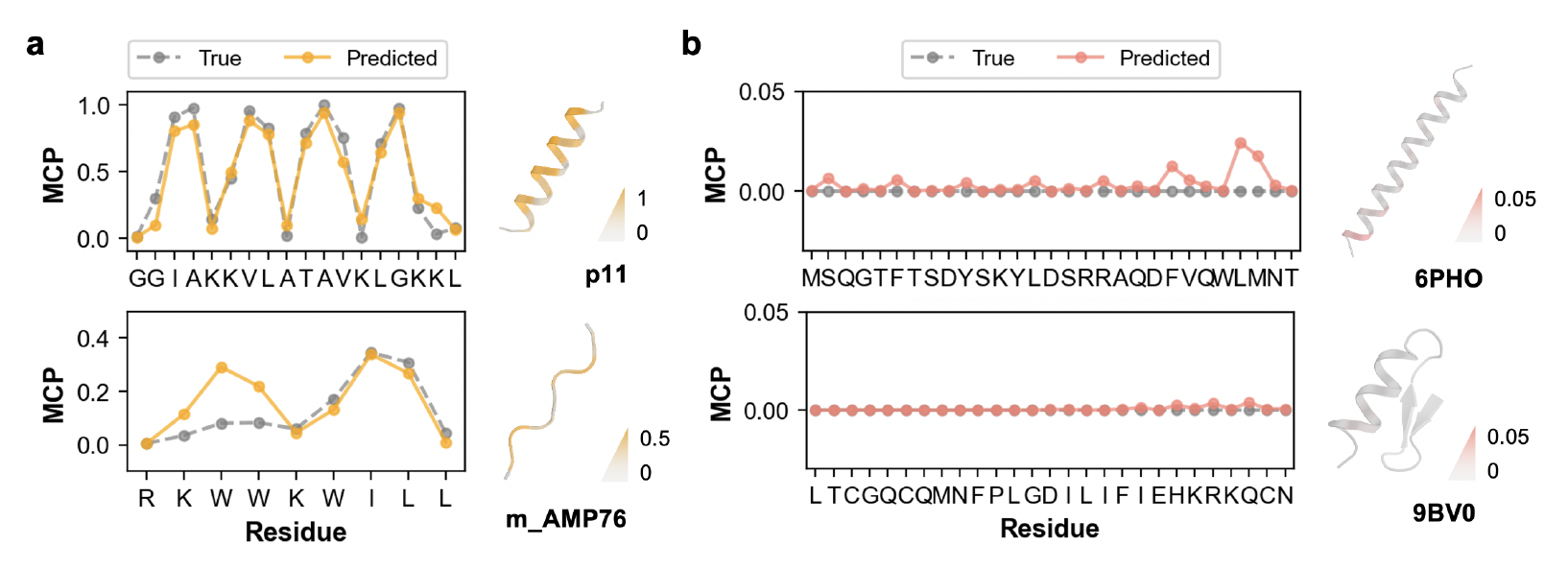
Case studies of PepMCP predictions. **a**. PepMCP predicted values, true MCP values from MD, and structures of two membrane-lytic AMPs. **b**. PepMCP predicted values, true MCP values with all zeros, and structures of two soluble non-AMPs. Structures were predicted using AlphaFold2 and colored with predicted MCP values. The maximum values of colormaps were set to 1.0, 0.5, 0.05, and 0.05, respectively, according to the line plots.

This indicates that the 4-hop edge encoding in PepMCP was effective in aggregating the features along the *α*-helix.

### MemAMPdb Database and PepMCP Server

Based on the results above, we developed MemAMPdb database and the web server of PepMCP predictor for convenient use. In the MemAMPdb database (http://www.songlab.cn/MemAMPdb/), all 550 membrane-lytic AMPs (516 in the training set and 34 in the external test set) have been uploaded, along with their sequences, references, mechanism validation methods, and origins. Users can easily search by their ID, name, or keywords in mechanism assays and origin descriptions.

At the PepMCP server (http://www.songlab.cn/PepMCP/Introduction/), users can submit their query peptide sequences in FASTA format. The per-residue PepMCP predictions and line plots for each peptide can be downloaded after running PepMCP on our cloud server. Batch prediction is supported for no more than 20 sequences.

## Conclusion

To overcome the limitations of previous MCP predictors for peptides, we developed PepMCP, a graph-based peptide-tailored MCP predictor. PepMCP was trained using residue-level MCP labels from MD simulations of membrane-lytic AMPs, with the membranes composed of POPE and POPG lipids. We employed GraphSAGE in PepMCP to inductively pass the node-level features from the ESM C language model, and used 4-hop edges that effectively incorporated the spatial features of peptides, especially for *α*-helix. PepMCP significantly outperformed both ProtRAP-LM and the base model on these membrane-lytic AMPs. On an external test set, PepMCP captured the MCP value patterns across peptides at the node level and could distinguish the membrane-lytic AMPs from other soluble peptides at the sequence level. Therefore, PepMCP is a useful tool for recognizing and investigating membrane-lytic AMPs.

There are still some limitations to PepMCP. First, it favors *α*-helix peptides due to the imbalanced training data. Not only do *β*-sheet and coil peptides account for a small proportion of the membrane-lytic AMPs, but their AlphaFold-predicted structures used in MD simulations also exhibit lower confidence levels (Fig. S3b). We observe that PepMCP can still handle some peptides with other secondary structures among the test cases, but the predicted results for these require attention. Additionally, we suggest using PepMCP in conjunction with other antimicrobial predictors to identify novel membrane-lytic AMPs, given the limited number of training sequences available for PepMCP. Although PepMCP enables the capture of patterns or binding modes of peptide-membrane interactions, as reflected by MCP values, it cannot distinguish some subtle effects caused by membrane type or morphology, nor can it elucidate the detailed modes of action of membrane-lytic AMPs, such as barrel-stave or carpet models.

PepMCP is trained only on membrane-lytic AMPs, but it has the potential to generalize to other membrane-active peptides. Examples include cell-penetrating peptides, which permeate the membrane, and transmembrane signal peptides. As shown in Fig. S4, PepMCP produces MCP predictions of around 0.5 for nearly all residues in these membrane-active *α*-helical peptides, except at the terminus. We expect that PepMCP will play a crucial role in the discovery of membrane-lytic AMPs and other membrane-active peptides in future studies.

## Author Contributions

C.S. conceived the project. R.D. developed the model and analyzed the results. T.A. conducted the MD simulations. Q.C. collected the data. K.K. and L.W. assisted in building the server and analyzed the results. Z.Z. assisted in processed the data. R.D., T.A., and Q.C. wrote the original manuscript, C.S. reviewed the manuscript. All authors have approved the final manuscript.

## Data Availability

The code and data are available at https://github.com/ComputBiophys/PepMCP. MemAMPdb is accessible at http://www.songlab.cn/MemAMPdb/. PepMCP server is accessible at http://www.songlab.cn/PepMCP/Introduction/.

## References

M. Abraham, T. Murtola, R. Schulz, S. Páll, J. Smith, B. Hess, and E. Lindahl. GROMACS: High performance molecular simulations through multi-level parallelism from laptops to supercomputers. SoftwareX, 1:19–25, 2015.

E. E. Ambroggio, F. Separovic, J. H. Bowie, G. D. Fidelio, and L. A. Bagatolli. Direct visualization of membrane leakage induced by the antibiotic peptides: Maculatin, citropin, and aurein. Biophysical Journal, 89(3):1874–1881, 2005.

H. G. Boman. Antibacterial peptides: basic facts and emerging concepts. Journal of Internal Medicine, 254(3):197–215, 2003.

A. K. Buck, D. E. Elmore, and L. E. Darling. Using fluorescence microscopy to shed light on the mechanisms of antimicrobial peptides. Future Medicinal Chemistry, 11(18):2447–2460, 2019.

G. Bussi, D. Donadio, and M. Parrinello. Canonical sampling through velocity rescaling. The Journal of Chemical Physics, 126(1):014101, 2007.

A. Chatzigoulas and Z. Cournia. Predicting protein–membrane interfaces of peripheral membrane proteins using ensemble machine learning. Briefings in Bioinformatics, 23(2): bbab518, 2022.

D. H. de Jong, G. Singh, W. F. D. Bennett, C. Arnarez, T. A. Wassenaar, L. V. Schäfer, X. Periole, D. P. Tieleman, and S. J. Marrink. Improved parameters for the Martini coarse-grained protein force field. Journal of Chemical Theory and Computation, 9(1):687–697, 2013.

R. Dong, Q. Cao, and C. Song. Painting peptides with antimicrobial potency through deep reinforcement learning. Advanced Science, 12(43):e06332, 2025.

J. Du, S. Zhang, G. Wu, J. M. F. Moura, and S. Kar. Topology adaptive graph convolutional networks. arXiv, 2018.

A. Elnaggar, H. Essam, W. Salah-Eldin, W. Moustafa, M. Elkerdawy, C. Rochereau, and B. Rost. Ankh: optimized protein language model unlocks general-purpose modelling. bioRxiv, page 2023.01.16.524265, 2023.

W. L. Hamilton, Z. Ying, and J. Leskovec. Inductive representation learning on large graphs. In NIPS, 2017.

Z. Han, L. Wang, and C. Song. Improved anisotropic network models for membrane and mechanosensitive ion channels. 2025.05.22.654704, 2025. protein dynamics bioRxiv, page

R. E. Harris, R. D. Whitehead III, and A. T. Alexandrescu. Solution structure of the Z0 domain from transcription repressor BCL11A sheds light on the sequence properties of protein-binding zinc fingers. Protein Science, 34(4):e70097, 2025.

T. Hayes, R. Rao, H. Akin, N. J. Sofroniew, D. Oktay, Z. Lin, R. Verkuil, V. Q. Tran, J. Deaton, M. Wiggert, R. Badkundri, I. Shafkat, J. Gong, A. Derry, R. S. Molina, N. Thomas, Y. A. Khan, C. Mishra, C. Kim, L. J. Bartie, M. Nemeth, P. D. Hsu, T. Sercu, S. Candido, and A. Rives. Simulating 500 million years of evolution with a language model. Science, 387(6736):850–858, 2025.

R. Heffernan, K. Paliwal, J. Lyons, J. Singh, Y. Yang, and Y. Zhou. Single-sequence-based prediction of protein secondary structures and solvent accessibility by deep whole-sequence learning. Journal of Computational Chemistry, 39 (26):2210–2216, 2018.

J. Huang, Y. Xu, Y. Xue, Y. Huang, X. Li, X. Chen, Y. Xu, D. Zhang, P. Zhang, J. Zhao, and J. Ji. Identification of potent antimicrobial peptides via a machine-learning pipeline that mines the entire space of peptide sequences. Nature Biomedical Engineering, 7(6):797–810, 2023.

S. Jain, J. Beazer, J. A. Ruffolo, A. Bhatnagar, and A. Madani. E1: Retrieval-augmented protein encoder models. bioRxiv, page 2025.11.12.688125, 2025.

J. Jumper, R. Evans, A. Pritzel, T. Green, M. Figurnov, O. Ronneberger, K. Tunyasuvunakool, R. Bates, A. Žídek, A. Potapenko, A. Bridgland, C. Meyer, S. A. A. Kohl, A. J. Ballard, A. Cowie, B. Romera-Paredes, S. Nikolov, R. Jain, J. Adler, T. Back, S. Petersen, D. Reiman, E. Clancy, M. Zielinski, M. Steinegger, M. Pacholska, T. Berghammer, S. Bodenstein, D. Silver, O. Vinyals, A. W. Senior, K. Kavukcuoglu, P. Kohli, and D. Hassabis. Highly accurate protein structure prediction with AlphaFold. Nature, 596(7873):583–589, 2021.

N. G. O. Júnior, C. M. Souza, D. F. Buccini, M. H. Cardoso, and O. L. Franco. Antimicrobial peptides: structure, functions and translational applications. Nature Reviews Microbiology, 23:687–700, 2025.

T. Kipf and M. Welling. Semi-supervised classification with graph convolutional networks. arXiv, abs/1609.02907, 2016.

J. Li, H. Guo, and C. Song. MemConverter: An iterative pipeline for reprogramming protein localization in membrane or aqueous solution. bioRxiv, page 2025.10.23.684164, 2025a.

J. Li, C. Yang, R. Dong, J. F. B. Juarez, L. Wang, M. E. Wettstein, D. Wang, C. Cao, Y. Lu, and C. Song. Mechanism-driven screening of membrane-targeting and pore-forming antimicrobial peptides. Advanced Science, page e16470, 2025b.

W. Li, L. Jaroszewski, and A. Godzik. Clustering of highly homologous sequences to reduce the size of large protein databases. Bioinformatics, 17(3):282–283, 2001.

W. Li, B. Huang, M. Guo, Z. Zeng, T. Cai, L. Feng, X. Zhang, L. Guo, X. Jiang, Y. Yin, E. Wang, X. Huang, and J. Zheng. Unveiling the evolution of antimicrobial peptides in gut microbes via foundation-model-powered framework. Cell Reports, 44(6):115773, 2025c.

A. L. Lomize, S. C. Todd, and I. D. Pogozheva. Spatial arrangement of proteins in planar and curved membranes by PPM 3.0. Protein Science, 31(1):209–220, 2022.

Y. Ma, Z. Guo, B. Xia, Y. Zhang, X. Liu, Y. Yu, N. Tang, X. Tong, M. Wang, X. Ye, J. Feng, Y. Chen, and J. Wang. Identification of antimicrobial peptides from the human gut microbiome using deep learning. Nature Biotechnology, 40: 1–11, 06 2022.

N. Michaud-Agrawal, E. J. Denning, T. B. Woolf, and O. Beckstein. MDAnalysis: A toolkit for the analysis of molecular dynamics simulations. Journal of Computational Chemistry, 32(10):2319–2327, 2011.

M. Mirdita, K. Schütze, Y. Moriwaki, L. Heo, S. Ovchinnikov, and M. Steinegger. ColabFold: making protein folding accessible to all. Nature Methods, 19:679–682, 2022.

L. Monticelli, S. K. Kandasamy, X. Periole, R. G. Larson, D. P. Tieleman, and S.-J. Marrink. The MARTINI coarse-grained force field: Extension to proteins. Journal of Chemical Theory and Computation, 4(5):819–834, 2008.

T. D. Newport, M. S. Sansom, and P. J. Stansfeld. The MemProtMD database: a resource for membrane-embedded protein structures and their lipid interactions. Nucleic Acids Research, 47(D1):D390–D397, 2018.

D. Paranou, A. Chatzigoulas, and Z. Cournia. Using deep learning and large protein language models to predict protein–membrane interfaces of peripheral membrane proteins. Bioinformatics Advances, 4(1):vbae078, 2024.

M. Parrinello and A. Rahman. Polymorphic transitions in single crystals: A new molecular dynamics method. Journal of Applied Physics, 52(12):7182–7190, 1981.

X. Periole, M. Cavalli, S.-J. Marrink, and M. A. Ceruso. Combining an elastic network with a coarse-grained molecular force field: Structure, dynamics, and intermolecular recognition. Journal of Chemical Theory and Computation, 5(9):2531–2543, 2009.

A. B. Poma, M. Cieplak, and P. E. Theodorakis. Combining the MARTINI and structure-based coarse-grained approaches for the molecular dynamics studies of conformational transitions in proteins. Journal of Chemical Theory and Computation, 13(3):1366–1374, 2017.

Y. Qi, H. I. Ingólfsson, X. Cheng, J. Lee, S. J. Marrink, and W. Im. CHARMM-GUI Martini Maker for coarse-grained simulations with the martini force field. Journal of Chemical Theory and Computation, 11(9):4486–4494, 2015.

C. D. Santos-Júnior, M. D. Torres, Y. Duan, Álvaro Rodríguez del Río, T. S. Schmidt, H. Chong, A. Fullam, M. Kuhn, C. Zhu, A. Houseman, J. Somborski, A. Vines, X.-M. Zhao, P. Bork, J. Huerta-Cepas, C. de la Fuente-Nunez, and L. P. Coelho. Discovery of antimicrobial peptides in the global microbiome with machine learning. Cell, 187(14): 3761–3778.e16, 2024.

F. Suarez-Leston, M. Calvelo, G. F. Tolufashe, A. Muñoz, U. Veleiro, C. Porto, M. Bastos, Añgel Piñeiro, and R. Garcia-Fandino. SuPepMem: A database of innate immune system peptides and their cell membrane interactions. Computational and Structural Biotechnology Journal, 20:874–881, 2022.

P. Szymczak, W. Zarzecki, J. Wang, Y. Duan, J. Wang, L. P. Coelho, C. de la Fuente-Nunez, and E. Szczurek. AI-driven antimicrobial peptide discovery: Mining and generation. Accounts of Chemical Research, 58(12):1831–1846, 2025.

N. van Hilten, N. Verwei, J. Methorst, C. Nase, A. Bernatavicius, and H. J. Risselada. PMIpred: a physicsinformed web server for quantitative protein–membrane interaction prediction. Bioinformatics, 40(2):btae069, 2024.

P. Veličković, G. Cucurull, A. Casanova, A. Romero, P. Lió, and Y. Bengio. Graph attention networks. arXiv, 2018.

L. Wang, J. Zhang, D. Wang, and C. Song. Membrane contact probability: An essential and predictive character for the structural and functional studies of membrane proteins. PLOS Computational Biology, 18(3):e1009972, 2022.

L. Wang, K. Kang, and C. Song. ProtRAP-LM: Fast and accurate protein relative accessibility prediction and membrane protein screening through protein language model embeddings. bioRxiv, page 2025.01.20.633985, 2025a.

M. Wang, D. Zheng, Z. Ye, Q. Gan, M. Li, X. Song, J. Zhou, C. Ma, L. Yu, Y. Gai, T. Xiao, T. He, G. Karypis, J. Li, and Z. Zhang. Deep graph library: A graph-centric, highly-performant package for graph neural networks. arXiv, 2019.

Y. Wang, L. Zhao, Z. Li, Y. Xi, Y. Pan, G.-P. Zhao, and L. Zhang. A generative artificial intelligence approach for the discovery of antimicrobial peptides against multidrugresistant bacteria. Nature Microbiology, 10:2997–3012, 2025b.

E. L. Wu, X. Cheng, S. Jo, H. Rui, K. C. Song, E. M. Dávila-Contreras, Y. Qi, J. Lee, V. Monje-Galvan, R. M. Venable,J. B. Klauda, and W. Im. CHARMM-GUI Membrane Builder toward realistic biological membrane simulations. J Comput Chem, 35(27):1997–2004, 2014.

