## Supplementary Information for "PepMCP: A Graph-Based Membrane Contact Probability Predictor for Membrane-Lytic Antimicrobial Peptides"

### Supplementary Figures

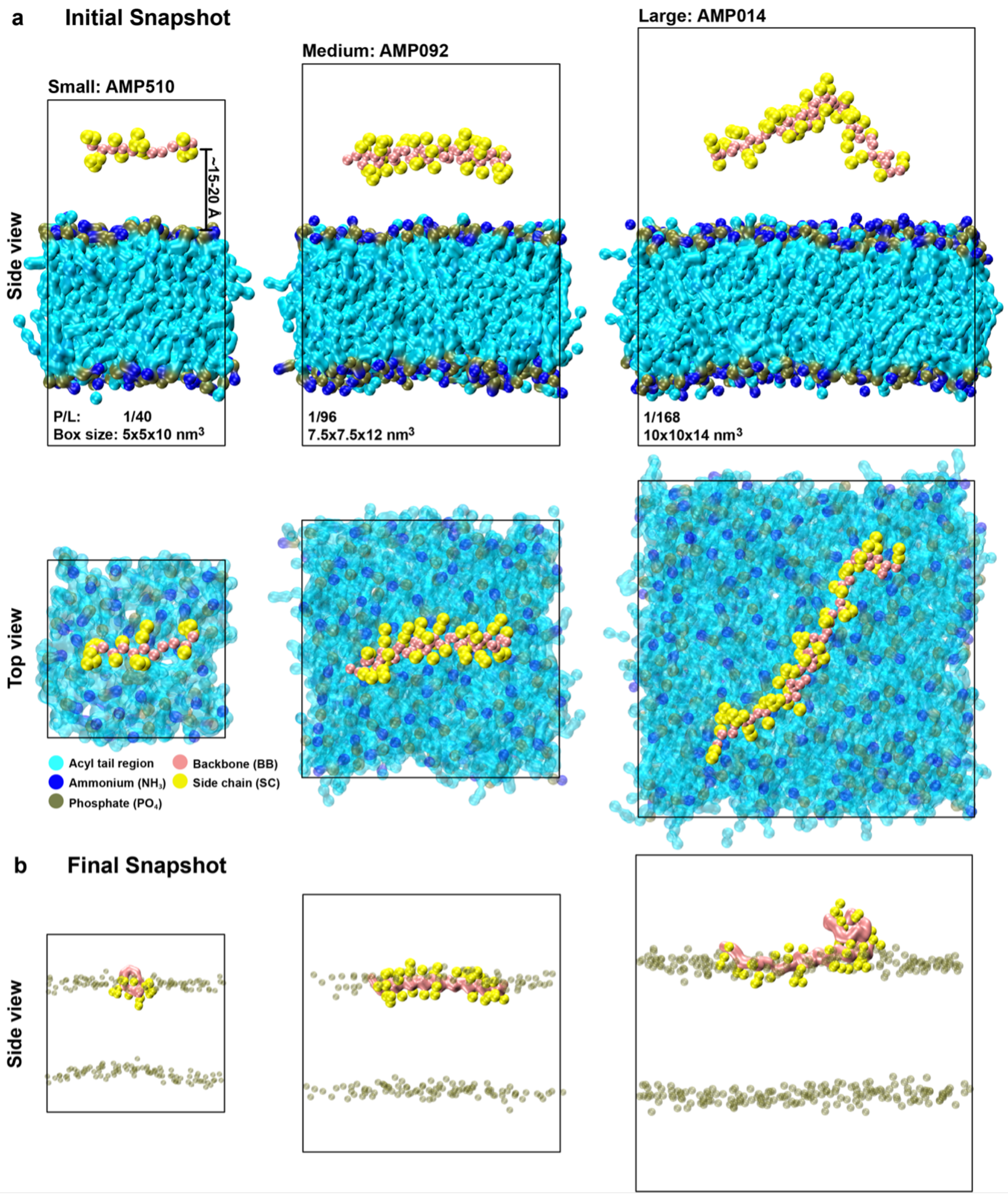


**Figure S1:** Coarse-grained (CG) simulation systems for peptide–membrane interactions. **a**. Initial side and top views of small, medium, and large simulation boxes showing the placement of peptides above the membrane, with the peptide center of mass positioned ∼15–20 Å from the membrane headgroup region. Corresponding peptide-to-lipid (P/L) ratios and box dimensions are indicated, along with representative peptide conformations within each simulation box. **b**. Final side view snapshots of each system. The cases for each system size were: small-AMP510 (synthetic peptide #14d), medium—AMP092 (MAP34-B), and large—AMP014 (prosthecin-1), respectively. The small and medium systems were at 2 µs, while the large was at 3 µs. Color scheme: lipid acyl tail region (cyan), ammonium headgroups (blue), phosphate headgroups (olive), peptide backbone (salmon), and side chains (yellow). Molecular structures were visualized using VMD version 1.9.4.

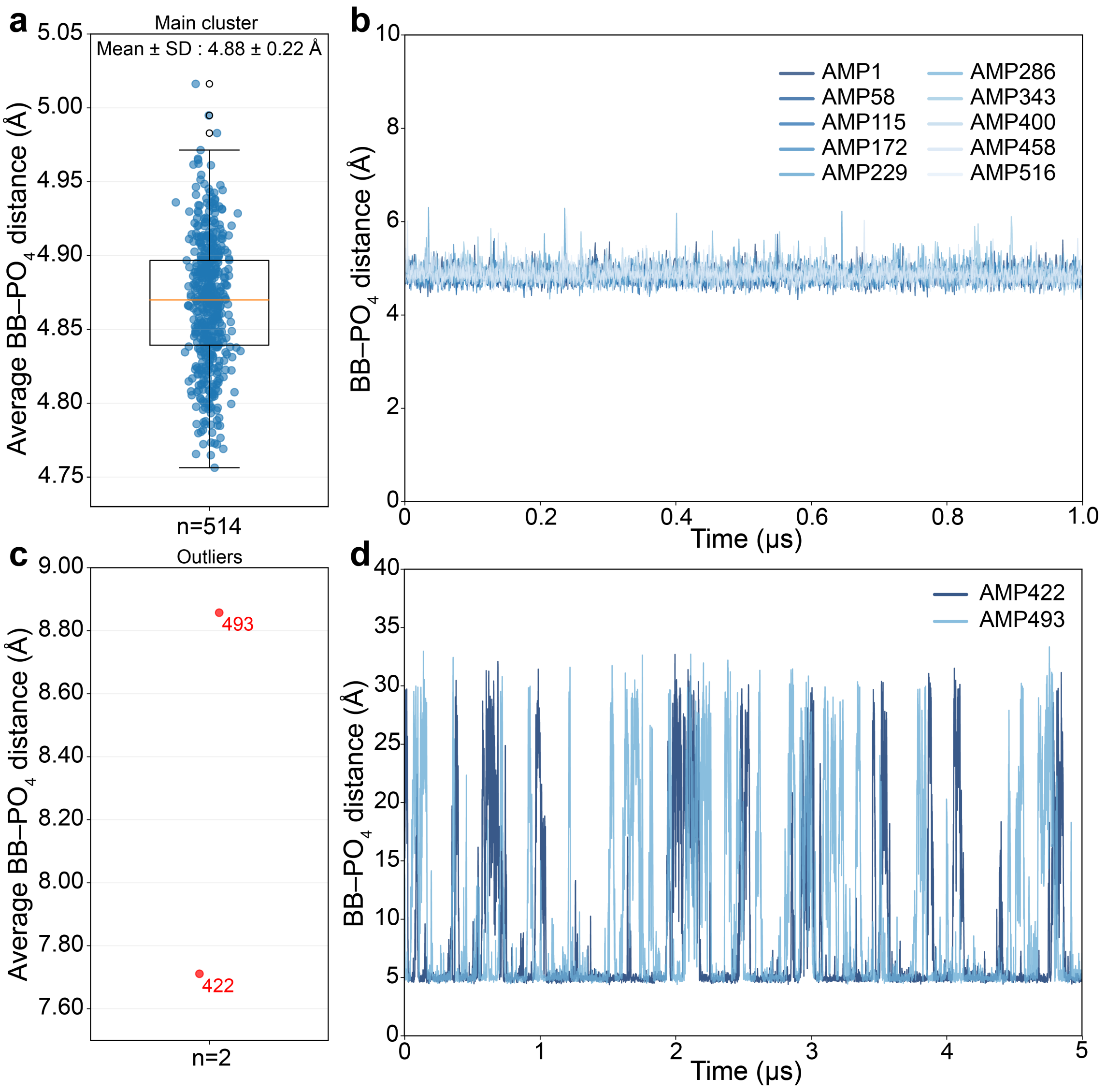


**Figure S2:** Binding of membrane-lytic AMPs with membranes in coarse-grained molecular dynamics simulations. The binding is characterized by the distribution and time evolution of backbone–phosphate (BB–$PO_4$) minimum distances, where BB represents the backbone beads of the AMPs, and $PO_4$ represents the phosphates of lipid molecules. **a**. Average BB–$PO_4$ minimum distances over the final 1 μs for 514 out of the 516 membrane-lytic AMP dataset. Each blue dot represents the time-averaged value for a single system. **b**. Time evolution of BB–$PO_4$ minimum distances for ten representative AMP systems during the final 1 μs, demonstrating that these peptides are stably bound to the membrane surface. **c**. Average BB–$PO_4$ minimum distances over the final 1 μs for the two outlier systems (AMP422 and AMP493), exhibiting significantly larger average BB–$PO_4$ distances. **d**. Time evolution of BB–$PO_4$ minimum distances for AMP422 and AMP493 over 5 μs trajectories, demonstrating that these two peptides could not stably bind to the membrane surface.

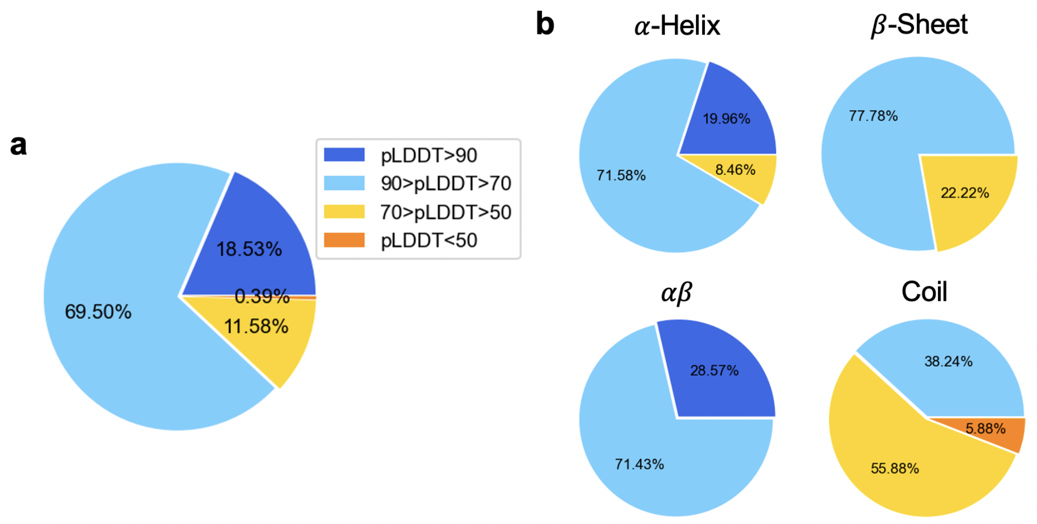


**Figure S3:** The confidence of 516 AlphaFold-predicted peptide structures. **a**. pLDDT distribution of 516 peptides. **b**. pLDDT distribution of peptides in different secondary structures.

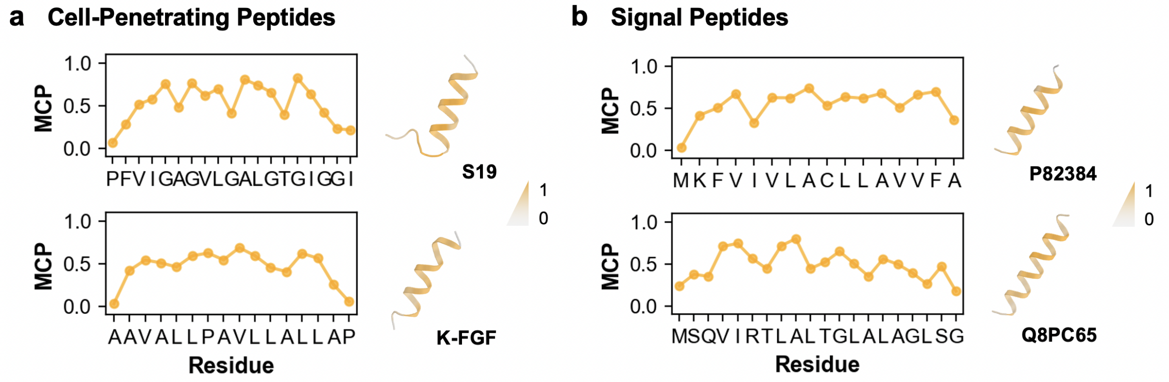


**Figure S4:** Case studies of PepMCP on other membrane-active peptides. **a**. PepMCP predicted values and structures of two cell-penetrating peptides. **b**. PepMCP predicted values and structures of two cell-penetrating peptides. Structures were predicted using AlphaFold2 and colored with predicted MCP values.
